# Establishing evidenced-based best practice for the *de novo* assembly and evaluation of transcriptomes from non-model organisms

**DOI:** 10.1101/035642

**Authors:** Matthew D. MacManes

## Abstract

Characterizing transcriptomes in both model and non-model organisms has resulted in a massive increase in our understanding of biological phenomena. This boon, largely made possible via high-throughput sequencing, means that studies of functional, evolutionary and population genomics are now being done by hundreds or even thousands of labs around the world. For many, these studies begin with a *de novo* transcriptome assembly, which is a technically complicated process involving several discrete steps. Each step may be accomplished in one of several different ways, using different software packages, each producing different results. This analytical complexity begs the question – *Which method(s) are optimal?* Using reference and non-reference based evaluative methods, I propose a set of guidelines that aim to standardize and facilitate the process of transcriptome assembly. These recommendations include the generation of between 20 million and 40 million sequencing reads from single individual where possible, error correction of reads, gentle quality trimming, assembly filtering using `Transrate` and/or gene expression, annotation using `dammit`, and appropriate reporting. These recommendations have been extensively benchmarked and applied to publicly available transcriptomes, resulting in improvements in both content and contiguity. To facilitate the implementation of the proposed standardized methods, I have released a set of version controlled open-sourced code, `The Oyster River Protocol for Transcriptome Assembly`, available at http://oyster-river-protocol.rtfd.org/.

## Introduction

For all biology, modern sequencing technologies has provided for an unprecedented opportunity to gain a deep understanding of genome level processes that underlie a very wide array of natural phenomenon, from intracellular metabolic processes to global patterns of population variability. Transcriptome sequencing has been influential, particularly in functional genomics, and has resulted in discoveries not possible even just a few years ago. This in large part is due to the scale at which these studies may be conducted. Unlike studies of adaptation based on one or a small number of candidate genes (e.g. (Fitzpatrick et al., 2005; Panhuis, 2006)), modern studies may assay the entire suite of expressed transcripts – the transcriptome – simultaneously. In addition to issues of scale, as a direct result of enhanced dynamic range, newer sequencing studies have increased ability to simultaneously reconstruct and quantitate lowly- and highly-expressed transcripts, (Vijay et al., 2013; Wolf, 2013). Lastly, improved methods for the detection of differences in gene expression (*e.g.,* (Love et al., 2014; Robinson et al., 2010)) across experimental treatments has resulted in increased resolution for studies aimed at understanding changes in gene expression.

As a direct result of their widespread popularity, a diverse toolset for the assembly and analysis of transcriptome exists. Notable amongst the wide array of tools include several for quality visualization - `FastQC` (available here) and `SolexaQA` (Cox et al., 2010), read trimming (e.g. `Skewer` (Jiang et al., 2014), `Trimmomatic` (Bolger et al., 2014) and `Cutadapt` (Martin, 2011)), read normalization (`khmer` (Pell et al., 2012)), error correction (Le et al., 2013), assembly (`Trinity` (Haas et al., 2013), `SOAPdenovoTrans` (Xie et al., 2014)), and assembly verification (`Transrate` (Smith-Unna et al., 2015)), `BUSCO` (<u>B</u>enchmarking <u>U</u>niversal <u>S</u>ingle-<u>C</u>opy <u>O</u>rthologs – (Simüo et al., 2015)), and `RSEM-eval` (Li et al., 2014)). The ease with which these tools may be used to produce transcriptome assemblies belies the true complexity underlying the overall process. Indeed, the subtle (and not so subtle) methodological challenges associated with transcriptome reconstruction may result in highly variable assembly quality. Amongst the most challenging include isoform reconstruction and simultaneous assembly of low- and high-coverage transcripts (Johnson et al., 2003; Modrek et al., 2001), which together make accurate transcriptome assembly technically challenging. Production of an accurate transcriptome assembly requires a large investment in time and resources. Each step in it’s production requires careful consideration. Here, I propose a set of evidence-based guidelines for assembly and evaluation that will result in the production of the highest quality transcriptome assembly possible.

Currently, a very large number of labs and research programs depend, often critically, on the production of accurate transcriptome resources. That no current best practices exits – particularly for those working in non-model systems – has resulted in an untenable situation where each laboratory makes up it’s own computational pipeline. These pipelines, often devoid of rigorous quality evaluation, may have important downstream consequences. This manuscript, by proposing a specific evidence-based process, significantly enhances the technical quality and reproducibility of transcriptome studies, which is critical for this emerging field of research.

## Methods

To demonstrate the merits of my recommendations, a large number of assemblies were produced using a variety of methods. For all assemblies performed, Illumina sequencing adapters were removed from both ends of the sequencing reads, as were nucleotides with quality Phred ≤ 2, using the program `Trimmomatic` version 0.32 (Bolger et al., 2014). The reads were assembled using Trinity release 2.1.1 (Haas et al., 2013) using default settings. `Trinity` was used as the default assembler as it has been previously reported to be best in class (Li et al., 2014; Smith-Unna et al., 2015). Assemblies were characterized using `Transrate` version 1.0.1 (Smith-Unna et al., 2015). Using this software, I generated three kinds of metrics: contig metrics; mapping metrics which used as input the same reads that were fed into the assembler for each assembly; and comparative metrics which used as input the *Mus musculus* version 75 transcriptome. In addition to the metrics provided by `Transrate`, I evaluated completeness of each assembly by use of `BUSCO`, a software package that searches for highly conserved, near-universal, single copy orthologs. All assemblies generated are available here, and will be moved to Dryad on acceptance.

To understand the influence of read depth on assembly quality, I produced subsets of size 1,2,5,10,20,40,60,80,100 million paired end reads of two publicly available paired-end datasets A *Mus* dataset -SRR797058 described in (Han et al., 2013) and a human dataset - SRR1659968. The subsampling procedure was accomplished via the software package seqtk (https://github.com/lh3/seqtk). For the evaluation of the effects of sequence polymorphism on assembly quality, I use reads from BioProject PRJNA157895 described in (MacManes and Lacey, 2012), a *Ctenomys* dataset which consists of 10 read files from the hypothalami of 10 different individuals. This dataset was assembled two ways. First, the reads from all 10 individuals were jointly assembled in one large assembly [CODE]. This assembly was compared to the assembly of a single individual [CODE]. Assemblies were generated and evaluated as per above.

To evaluate the effects of error correction, I used the subsampled read datasets, which were subsequently error corrected using the following software packages: `SEECER` version 0.1.3 (Le et al., 2013), `Lighter` version 1.0.7 (Song et al., 2014), `SGA` version 0.10.13 (Simpson and Durbin, 2012), `bfc` version r177 (Li, 2015), `RCorrector` (Song and Florea, 2015), and `BLESS` version 0.24 (Heo et al., 2014). In correction algorithms (`SGA, BLESS, bfc`) that allowed for the use of larger *kmer* lengths, I elected to error correct with a small (*k* = 31) and a long (*k* = 55) *kmer*, while for the other software (`RCorrector, SEECER` and `Lighter`) that does not allow for longer *kmer* values, I set *k* = 31. bfc requires interleaved reads, which was accomplished using khmer version 2.0 (Alameldin et al., 2015; Brown et al., 2012; McDonald and Brown, 2013). Code for performing these steps is available [here].

The effects of `khmer` digital normalization (Pell et al., 2012) were characterized by generating three 20 million and three 100 million read subsets of the larger *Mus* dataset. Digital normalization was performed using a median kmer abundance threshold of 30. The resulting datasets were assembled using `Trinity`, and evaluated using `BUSCO` and `Transrate`. Code for performing these steps is available in the `diginorm` target of the [Makefile].

Post-assembly processing was evaluated using several assembly datasets of various sizes, generated above. Each assembly was evaluated using `Transrate. Transrate` produces a score based on contig and mapping metrics, as well as a more optimal assembly where poorly supported contigs (putative assembly artifacts) are removed. Both the original and `Transrate` optimal assembly are evaluated using `BUSCO`, to help better understand if filtration results in the loss of non-artifactual transcripts. In addition to `Transrate` filtration, an additional, or alternative filtration step is performed using estimates of gene expression (TPM=transcripts per million). TPM is estimated by two different software packages that implement two distinct methods - `Salmon` (Patro et al., 2015) and `Kallisto` (Bray et al., 2015). Transcripts whose expression is estimated to be greater than a given threshold, typically TPM=1 or TPM=0.5 are retained. As above, the filtered assemblies are evaluated using `BUSCO`, to help better understand if filtration results in the loss of non-artifactual transcripts. Code for performing these steps is available in the `QC` target of the makefile available [here].

## Recommendations

### 0.1 Input Data

#### Summary Statement: Sequence 1 or more tissues from 1 individual to a depth of between 20 million and 40 million 100bp or longer paired-end reads

When planning to construct a transcriptome, the first question to ponder is the type and quantity of data required. While this will be somewhat determined by the specific goals of the study and availability of tissues, there are some general guiding principals. As of 2014, Illumina continues to offer the most flexibility in terms of throughout, analytical tractability, and cost (Glenn, 2011) and as a result, the recommendations here are primarily related to assembly using Illumina data. It is worth noting however, that long-read (e.g. PacBio) transcriptome sequencing is just beginning to emerge as an alternative (Au et al., 2013), particularly for researchers interested in understanding isoform complexity. Though currently lacking the throughput for accurate quantitation of gene expression, long read technologies, much like they have done for *de novo* genome assembly, seem likely to replace short-read-based *de novo* transcriptome assembly at some point in the future.

For the typical transcriptome study, one should plan to generate a reference based on 1 or more tissue types, with each tissue adding unique tissue-specific transcripts and isoforms. Though increasing the amount of sequence data collected does increase the accuracy and completeness of the assembly (Figure 1, 3) albeit marginally, a balance between cost and quality exists. For the datasets examined here (vertebrate tissues), sequencing more than between 20M and 40M paired-end reads is associated with the discovery of very few additional transcripts, and only minor improvement in other assembly metrics. Read length should be at least 100bp, with longer reads likely aiding in isoform reconstruction and contiguity (Garber et al., 2011). In the case where multiple tissues are sequenced, it is likely best to combine reads from each tissue together to produce a joint assembly.

Because sequence polymorphism increases the complexity of the *de bruijn* graph (Iqbal et al., 2012; Studholme, 2010), and therefore may negatively effect the assembly itself, the reference transcriptome should be generated from reads corresponding to as homogeneous a sample as possible. For outbred, non-model organisms, this usually means generating reads from a single individual. When more then one individual is required to meet other requirements (*e.g.*, for differential expression replicates or experimental treatment conditions), keeping the number of individuals to a minimum is paramount. For instance, when performing an experiment where a distinct set of genes may be expressed in different treatments (or sexes), the recommendation is to sequence one individual from each treatment class.

To illustrate this effect, I examined the effects of assembling reads from 10 individuals jointly, versus assembling a representative individual. This individual was selected based on having the highest number of reads. Using 30 threads on a standard Ubuntu workstation, the individual assembly of 38 million paired end read took approximately 23 hours and 20Gb of RAM, while the joint assembly took five days and 150Gb of RAM. In addition to this, to eliminate the potential confounding factor of increased coverage, I assembled another dataset that consisted of a random subsample of 3.8M paired-end reads from each of the 10 samples. This subsampled assembly used similar resources as did the single-individual assembly. Per Table 1, the joint assembly used more than eight times more reads, and is more than four times larger than the assembly of a single individual. Despite the additional read data, the `Transrate` score is markedly decreased, although the BUSCO statistics are slightly better. The large joint assembly suffers from major structural problems that are unfixable via the proposed filtering procedures. Specifically, read-mapping data suggests that 28.7% of the contigs in the joint assembly and 18% in the subsampled assembly could be merged, versus 15% in the single assembly. This structural problem is likely the result of sequence polymorphism and may cause significant issues for many common downstream processes.

**Table 1.** A comparison of the raw assemblies resulting from a single individuals versus the joint assembly of 10 individuals as well as a joint but subsampled dataset consisting of a random 3.8M reads from each of the 10 samples. The individual assembly of 38 million reads resulted in a high quality assembly as evidenced by the `Transrate` score of 0.3064 (per (Smith-Unna et al., 2015), this is a score better that 50% of published transcriptomes), and a high BUSCO score. The assembly of 10 individuals scores lower using `Transrate`, though a small number of new transcripts are discovered. The joint-subsampled assembly recovered an equivalent number of transcripts, but resulted in a lower `Transrate` score, indicating a lower quality assembly.

| Name | Num. Reads | Num. Contigs | Assembly Size | Score | BUSCO |
| --- | --- | --- | --- | --- | --- |
| **Single Ind.** | 38M | 205812 | 131.6Mb | 0.3064 | C:81%,D:41%,M:9% |
| **Subsampled** | 38M | 304162 | 183.8Mb | 0.2619 | C:84%,D:47%,M:8.4% |
| **10 Ind.** | 269M | 913295 | 440.2Mb | 0.22011 | C:88%,D:51%,M:5% |

**Figure 1.**
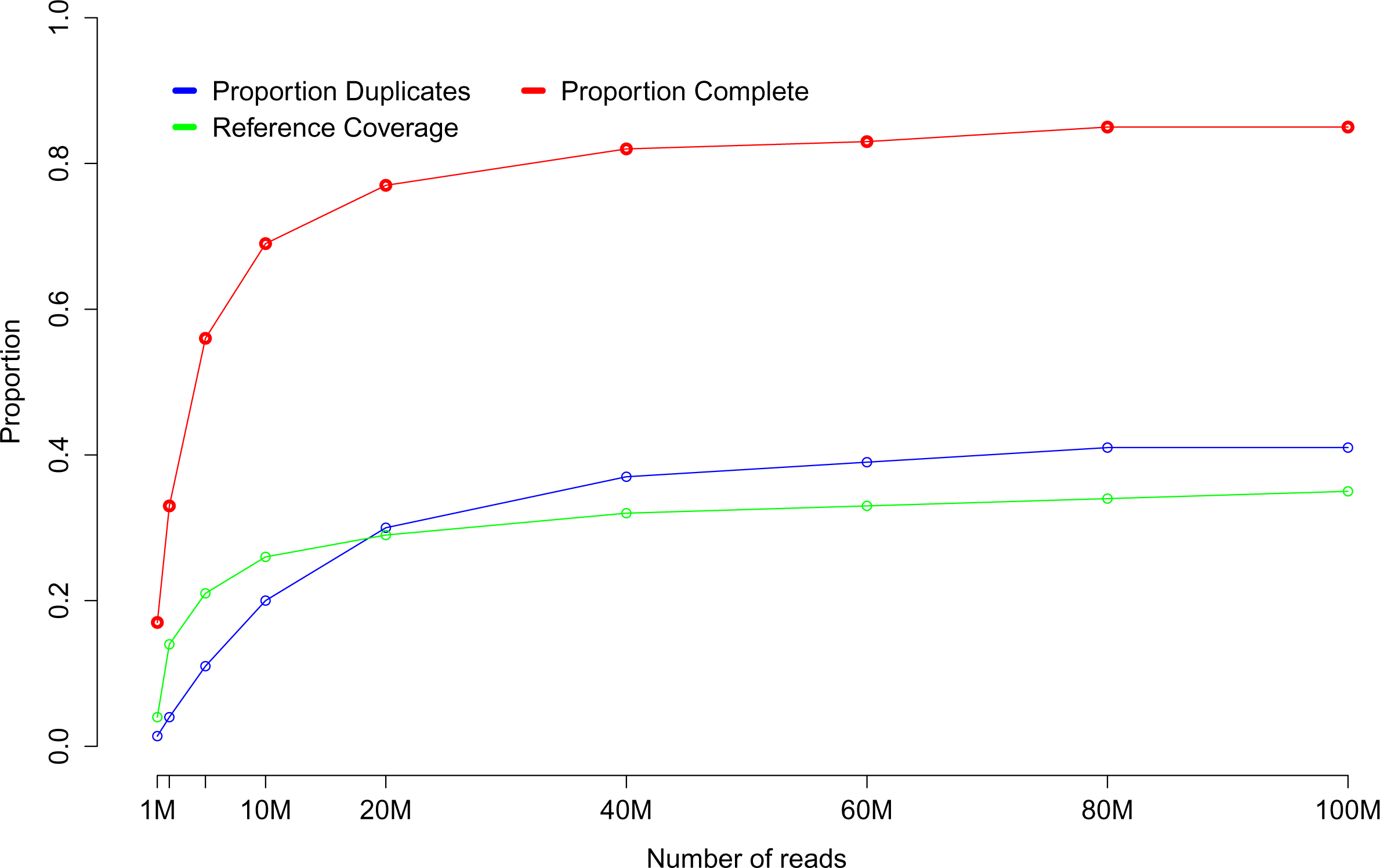
Assembly of multiple subsetted datasets suggests that sequencing beyond 20-40 million paired end reads does not results in further sequence discovery. Proportion complete indicates the proportion of BUSCOs that were found to be full length. Proportion duplicates are those BUSCOs that were found multiple times in the assembly dataset. Reference coverage is a `Transrate` generated metric indicating the proportion of the reference *Mus* transcriptome found in the *de novo* assembly. Higher numbers for reference coverage and proportion complete indicate a more complete assembly.

### 0.2 Quality Control of Sequence Read Data

#### Summary Statement:Visualize your read data. Error correct reads using bfc for low to moderately sized datasets and RCorrector for higher coverage datasets. Remove adapters, and employ gentle quality filtering using PHRED ≤ 2 as a threshold

Before assembly, it is critical that appropriate quality control steps are implemented. It is often helpful to generate some metrics of read quality on the raw data. Several software packages are available – I am fond of `SolexaQA` (Cox et al., 2010) and `FastQC`. Immediately upon download of the read dataset from the sequence provider, metrics of read quality, generated by either of these two software packages, should be generated. Of note – a copy of the raw reads should be compressed and archived, preferably on a physically separated device for long term archival storage. For this, I have successfully used Amazon S3 cloud storage, though many options exist.

Immediately after visualizing the raw data, error correction of the sequencing reads should be done (MacManes and Eisen, 2013). A very large number of read correction software packages exist, and several of them are benchmarked here using the *Mus* (Figure 2, and Tables S1-S11) and *Homo* datasets (Tables S12-S21). In all evaluated datasets, the error correction bfc was the best when correcting less than approximately 20M paired-end reads. When correcting more, the software `RCorrector` provided the optimal correction. The effects of error correction on assembly were evaluated using `BUSCO` and `Transrate`. While error correction did not result in significant improvements in `BUSCO` metrics, the transrate scores were substantially improved (Figure 3). These scores were largely improved by the fact that assemblies using error corrected reads had fewer low-covered bases and contigs, and a slightly higher mapping rate.

The error corrected reads are then subjected to vigorous adapter sequence removal, typically using `Trimmomatic` (Bolger et al., 2014) or `Skewer` (Jiang et al., 2014). With adapter sequence removal may be a quality trimming step. Here, substantial caution is required, as aggressive trimming has detrimental effects on assembly quality. Specifically, I recommend trimming at Phred=2 (MacManes, 2014), a threshold associated with removal of only the lowest quality bases. After adapter removal and quality trimming, the previously error corrected reads are now ready for *de novo* transcriptome assembly.

**Figure 2.**
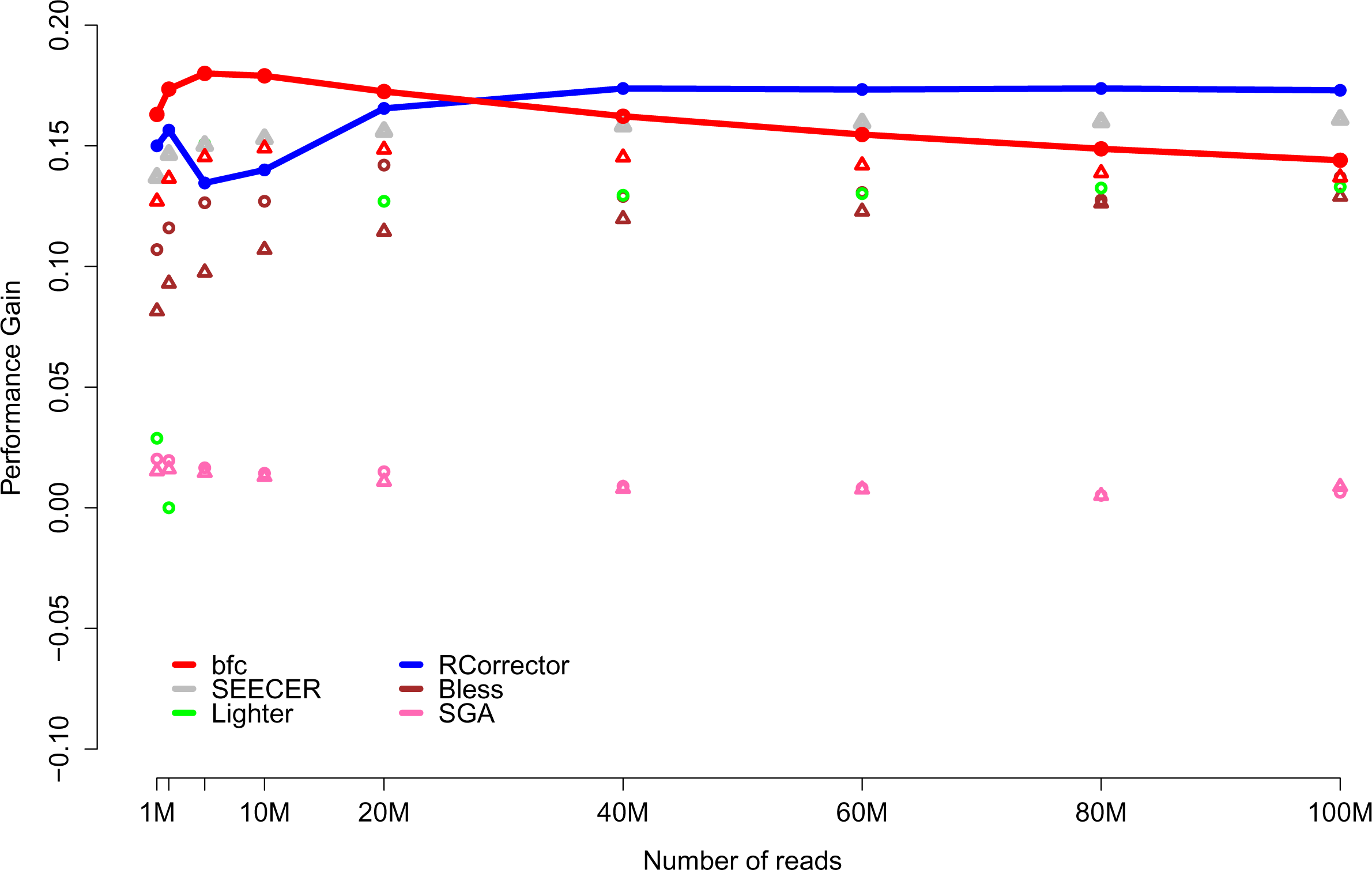
Error correction of reads results in a performance gain (defined as: (perfect error corrected reads – perfect raw reads) + reads made better – reads made worse). Perfect reads are reads that map to the reference without mismatch. Better and worse reads are those that map with fewer or more mismatches. Low coverage datasets are best corrected with bfc, which higher coverage datasets are optimally corrected with `RCorrector`. The best performing corrections improve the quality of more than 15% of reads. To emphasize the patterns of performance of the two best-performing error correctors, their points are connected by lines.

**Figure 3.**
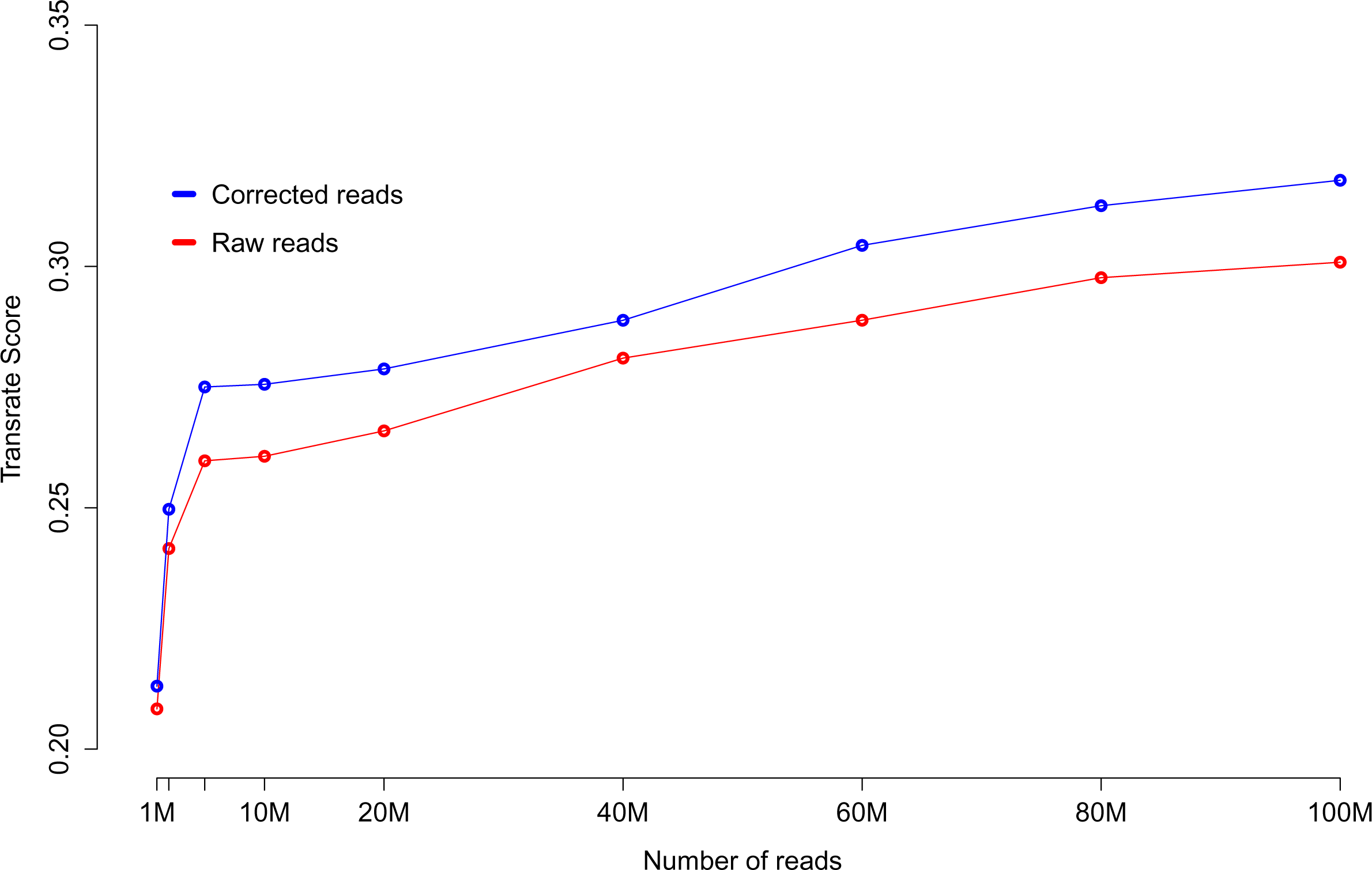
Error correction (with the best performing correction software, described in Figure 2), results in a consistent increase in the `Transrate` score, which indicates a higher quality assembly across all coverage depths.

### 0.3 Coverage normalization

#### Summary Statement: Normalize your data, only if you have to

Depending on the volume of input data, the availability of a high-memory workstation, and the rapidity with which the assembly is needed, coverage normalization may be employed. This process, which, using a streaming algorithm and measurement of the median kmer abundance of each read, aims to erode areas of high coverage while leaving untouched, reads spanning lower coverage areas. Normalization may be accomplished in the software package `khmer` (Pell et al., 2012), or within `Trinity` using a computational algorithm based on `khmer`. To evaluate khmer normalization, I generated 6 datasets, 3 using 20M paired reads and 3 using 100M paired reads. These datasets were assembled (available here) using Trinity and were evaluated via standard methods. These tests revealed that normalization did dramatically reduce RAM requirements and runtime, though it also decreased the number of complete BUSCO's found by an average of 1% for all assemblies. Normalization also decreased the `Transrate` score on average by 0.0248 for the 20M read assemblies and 0.0272 for the 100M read assemblies. Interestingly, normalization *increased* the percent BUSCO duplication by an average of only 1% for the smaller assembly, but by over 14% for the larger assembly. Given this, our recommendation is to employ digital normalization when the assembly is otherwise impossible, or when results are urgently needed, but that it should not be used by default for the production of transcriptome assemblies.

### 0.4 Assembly

#### Summary Statement: Assemble your data using Trinity, then remove poorly supported contigs

For non-model organisms lacking reference genomic resources, the error correction, adapter and quality trimming reads should be assembled *de novo* into transcripts. Currently, the assembly package `Trinity` (Haas et al., 2013) is thought to currently be the most accurate (Li et al., 2014), and therefore is recommended over other assemblers. While attempting a merged assembly with multiple assemblers may *ultimately* result in the highest quality assembly, options for merging assemblies are currently limited, and therefore is not recommended.

Trinity's underlying algorithm has been pre-optimized to recover large numbers of alternative isoforms, including many that are minimally supported by read data. As a result, in many cases, the raw assembly will require filtration to remove these assembly artifacts. Reference dependent and independent evaluative tools (*e.g.*, `Transrate, BUSCO`) allow for evidence-based post-assembly filtration. Typically, an initial quality-evaluation and filtration step is implemented using Transrate. This process assigns a score to the assembly, and creates an alternative assembly by removing contigs based on read-mapping metrics. This filtration step may result in the removal of a large proportion (as much as 67%) of the transcripts. Reference-based metrics are generated before and after this filtration step to ensure that filtration has not been too aggressive – that a significant number of known transcripts have not been removed. After `Transrate` filtration, or alternative to it, it is often helpful to employ a filtration step based on TPM. Because underlying assumptions of gene expression estimation software vary, which may result in variation of the actual estimates, gene expression is typically estimated using two different packages, `Salmon` and `Kallisto`. Transcripts whose abundance is less than either 1 or 0.5 are removed. Again, reference-based metrics are generated to ensure that a significant number of known transcripts are not removed.

The results of filtration on several datasets of varying size are presented in Figure 4. The reads used in the 1M,5M,10M,20M subset assemblies were corrected with `bfc`, while the reads for the larger assemblies were corrected with `RCorrector`. Each dataset was trimmed to a quality of Phred <2, and assembled with `Trinity`. The raw assembly was filtered by `Transrate` and by gene expression. `BUSCO` evaluation was performed before and after these filtration steps. In general, for low coverage datasets (less than 20 million reads), filtering based on expression, using TPM=1 as a threshold performs well, with `Transrate` filtering being too aggressive. With higher coverage data (more than 60 million reads) `Transrate` filtering may be optimal, as may gene expression filtering using a threshold of TPM=0.5. Again, the strength of this process is that it is guided by evidence, with filtering thresholds chosen based upon objective metrics.

**Figure 4.**
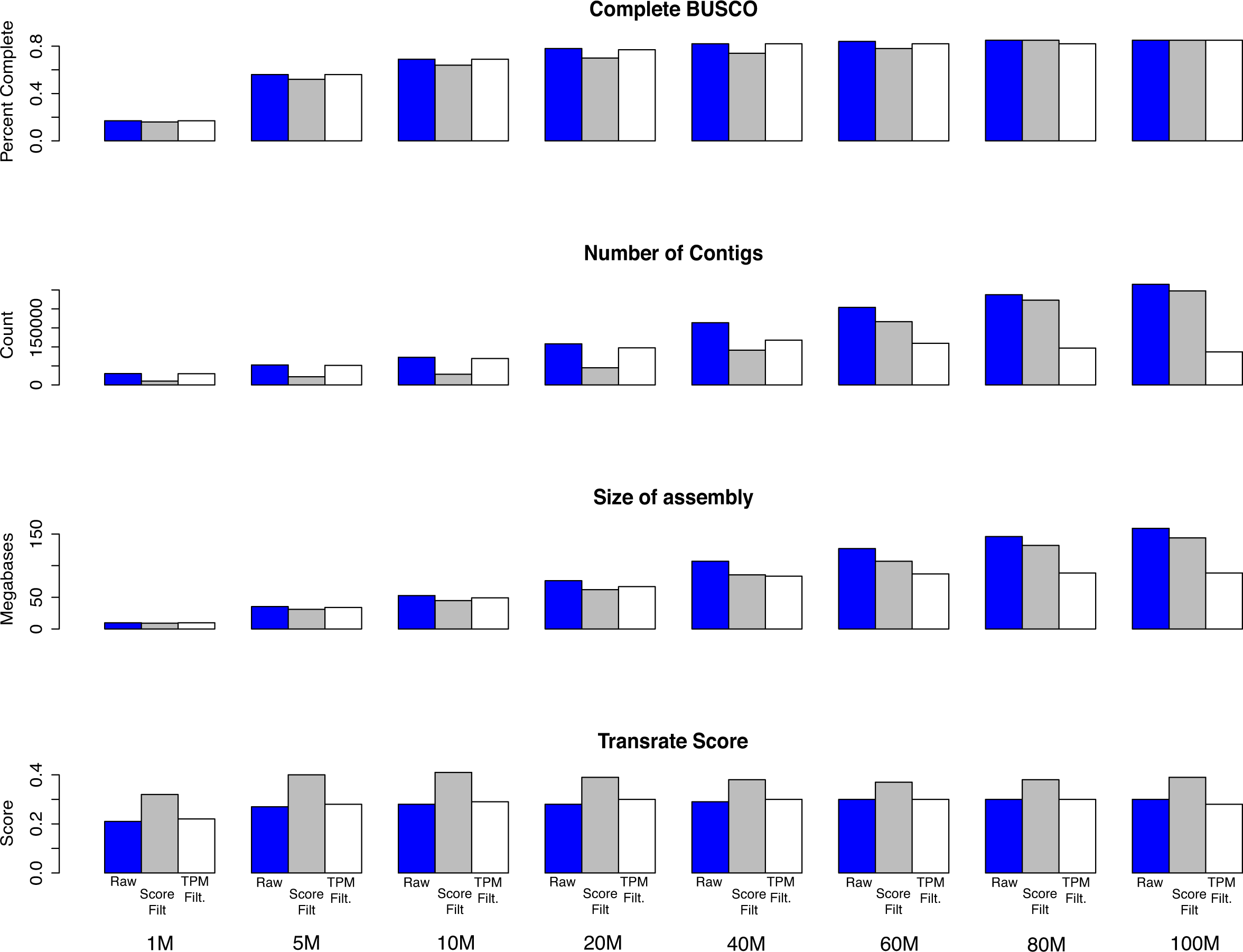
Post-assembly filtration. Using assemblies from the 1M,5M,10M,20M,40M,60M,80M,100M read subsets, I evaluated the effects of `Transrate` and TPM filtration using a threshold of TPM=1. Both `Transrate` and TPM filtering reduced the number of contigs and assembly size, though the magnitudes were dependent on the depth of sequencing. `BUSCO` scores were either decreased in some cases, or stable in others, representing the differential effects of filtering on different sized assemblies. In general, for low coverage datasets (less than 20 million reads), filtering based on expression, using TPM=1 as a threshold performs well, with `Transrate` filtering being too aggressive. With higher coverage data (more than 60 million reads) `Transrate` filtering may perform better, as mat expression filtering with a lower threshold.

### 0.5 Annotation, post-assembly quality verification, & reporting

#### Summary Statement: Verify the quality of your assembly using content based metrics. Annotate using dammit Report Transrate score, BUSCO statistics, number of unique transcripts, etc. Do not report meaningless statistics such as N50

Annotation is a critically important step in transcriptome assembly. Much like other steps, numerous options exist. Though the research requirements may drive the annotation process, I propose that a core set of annotations be provided with all *de novo* transcriptome assembly projects. The process through which these core annotations are accomplished is coordinated by the software package `dammit`. This software takes as input a fasta file and outputs a standard gff3 containing annotations. After annotation, but before downstream use, it is important to assess the quality of a transcriptome. Many authors have attempted to use typical genome assembly quality metrics for this purpose. In particular, N50 and other length-based summary statistic are often reported (e.g. (Hiz et al., 2014; Liang et al., 2013; Shinzato et al., 2014)). However, in addition to being a poor proxy for quality in genome assembly (Bradnam et al., 2013), N50 in the context of a transcriptome assembly carries very little information because the optimal contig length is not known (Li et al., 2014) – real transcripts vary greatly in length, ranging from tens of nucleotides to tens of thousands of nucleotides. Reportable metrics should be chosen based on their relevance for assembly optimization given the biological question at hand. In most cases, this means maximizing the number of transcripts that can be confidently attributed to the organism, while minimizing the number of technical artifacts related to the process of sequencing, quality control, and assembly. For many researchers, this means evaluation with both BUSCO and `Transrate`. The statistics found in Table 1 should be presented for all assemblies, with additional information supplementing these core vital statistics as needed.

## Testing the Oyster River Protocol

To evaluate the Oyster River Protocol for Transcriptome Assembly, I selected three publicly available Illumina RNAseq datasets and their corresponding assembled transcriptomes. These three assemblies included the Nile Tilapia, *Oreochromis niloticus* ((Zhang et al., 2013), SRR797490), an unpublished study of the Mediterranean black widow, *Latrodectus tredecimguttatus* (SRR954929), and lastly a work on *Delia antiqua* ((Guo et al., 2015), SRR916227). I analyzed the original transcriptomes using both `BUSCO` and `Transrate`, then followed the protocol as described here. Code for data analysis of the *Oreochromis* is available here. The other samples were processed in an identical fashion. The application of the Oyster River Protocol on these datasets resulted universally in a substantial (as much as 22%) improvement in the completeness of assemblies. Given a major goal of these types of studies includes reconstruction all expressed genes, this improvement may have substantial improvement on downstream work. The `Transrate` score was dramatically improved as well, particularly in the *Oreochromis* and *Delia* assemblies. This improvement speaks to the improvement of the structure of the assembly.

The filtering process through which these more optimal assemblies were is key. Evaluating both the BUSCO and `Transrate` scores before and after, allows for an objective way to decide if filtering has been too restrictive or not. Indeed, for the *Latrodectus* assembly, both `Transrate` and TPM filtering reduced the BUSCO score, while substantially increasing the `Transrate` score. Depending on the goals of the experiment, it may be determined that the structural integrity of the assembly outweighs improved content. In contrast to how post-assembly filtering is typically done, this method allow for the researcher to make an informed decision about these processes.

**Table 2.** The results of the application of the Oyster River Protocol to three available transcriptomes. Within each column, the 5 metrics, separated by forward slashes are: 1. The original assembly 2. The raw Trinity assembly 3. The `Transrate` filtered assembly 4. The TPM=1 filtered assembly, and 5. The `Transrate` filtered assembly that has been further filtered by expression. In all cases the assembly content, as evaluated by the `BUSCO` score is dramatically improved over the original assembly. These content-improved assemblies have acceptable `Transrate` scores, which in 2 of 3 cases are vastly superior to the scores of the original assembly.

| Name | Number Reads | Number Contigs | Assembly Size (Mb) | Transrate Score | BUSCO Score |
| --- | --- | --- | --- | --- | --- |
| **Oreochromis** | 25.2M | 79198/140035/100376/116038/88456 | 32.0/75.1/69.5/58.6/57.7 | 0.1103/0.2173/0.4778/0.2595/0.4479 | C:39%,M:46%/C:58%,M:28%/C |
| **Latrodectus** | 27.6M | 10259/36394/30932/27973/NA | 10.6/13.5/13.1/10.9/NA | 0.43673/0.2795/0.4968/0.338/NA | C:48%,M:38%/C:58%,M:28%/C |
| **Delia** | 25.8M | 29451/49099/38614/46145/32689 | 12.4/19.3/18.8/17.9/15.8 | 0.393/0.2036/0.4572/0.2305/0.4341 | C:40%,M:48%/C:62%,M:21%/C |

## Conclusions

With the rapid adoption of high-throughput sequencing, studies of functional, evolutionary and population genomics are now being done by hundreds or even thousands of labs around the world. These studies typically begin with a *de novo* transcriptome assembly. Assembly may be accomplished in one of several different ways, using different software packages, with each method producing different results. This complexity begs the question – *Which method(s) are optimal?* Using reference and non-reference based evaluative methods, I have proposed a set of guidelines `The Oyster River Protocol for Transcriptome Assembly` that aim to standardize and facilitate the process of transcriptome assembly. These recommendations include limiting assembly to between 20 million and 40 million sequencing reads from single individual where possible, error correction of reads, gently quality trimming, assembly filtering using `Transrate` or gene expression, annotation using `dammit`, and appropriate reporting. The processes result in a high quality transcriptome assembly appropriate for downstream usage.

## Acknowledgments

This work was significantly improved by discussions with Richard Smith-Unna, Brian Haas and many others. More generally, the work and it's presentation has been influenced by supporters of the Open Access and Science movements.

